# The DNA-based global positioning system—a theoretical framework for large-scale spatial genomics

**DOI:** 10.1101/2022.03.22.485380

**Authors:** Laura Greenstreet, Anton Afanassiev, Yusuke Kijima, Matthieu Heitz, Soh Ishiguro, Samuel King, Nozomu Yachie, Geoffrey Schiebinger

## Abstract

We present GPS-seq, a theoretical framework that enables massively scalable, optics-free spatial transcriptomics. GPS-seq combines data from high-throughput sequencing with manifold learning to obtain the spatial transcriptomic landscape of a given tissue section without optical microscopy. In this framework, similar to technologies like Slide-seq and 10X Visium, tissue samples are stamped on a surface of randomly-distributed DNA-barcoded spots (or beads). The transcriptomic sequences of proximal cells are fused to DNA barcodes, enabling the recovery of a transcriptomic pixel image by high-throughput sequencing. The barcode spots serve as “anchors” which also capture spatially diffused “satellite” barcodes, and therefore allow computational reconstruction of spot positions without optical sequencing or depositing barcodes to pre-specified positions. In theory, it could generate 100 mm × 100 mm spatial transcriptomic images with 10-20 μm resolution by localizing 10^8^ DNA-barcoded pixels with a single Illumina NovaSeq run. The general framework of GPS-seq is also compatible with standard single-cell (or single-nucleus) capture methods, and any modality of single-cell genomics, such as sci-ATAC-seq, could be transformed into spatial genomics in this strategy. We envision that GPS-seq will lead to breakthrough discoveries in diverse areas of biology by enabling organ-scale imaging of multiple genomic statuses at single-cell resolution for the first time.

## Introduction

Understanding the cellular architecture of tissues is a major challenge, with tremendous potential for impact across diverse areas of biology and medicine. Single-cell (sc)RNA-sequencing technologies enable high-dimensional, unbiased measurements of transcription levels across all genes in large populations of cells, but they lose the spatial context of cells within tissues^11-13^. Pioneering spatial transcriptomics (ST) technologies have recently made it possible to chart the spatial distribution of transcription profiles^14^. However, current ST technologies are limited in either resolution, or field of view (Table 1) and therefore cannot be used to analyze patterns of transcription over whole organs or organisms^1,3-7^.

**Table 1.**
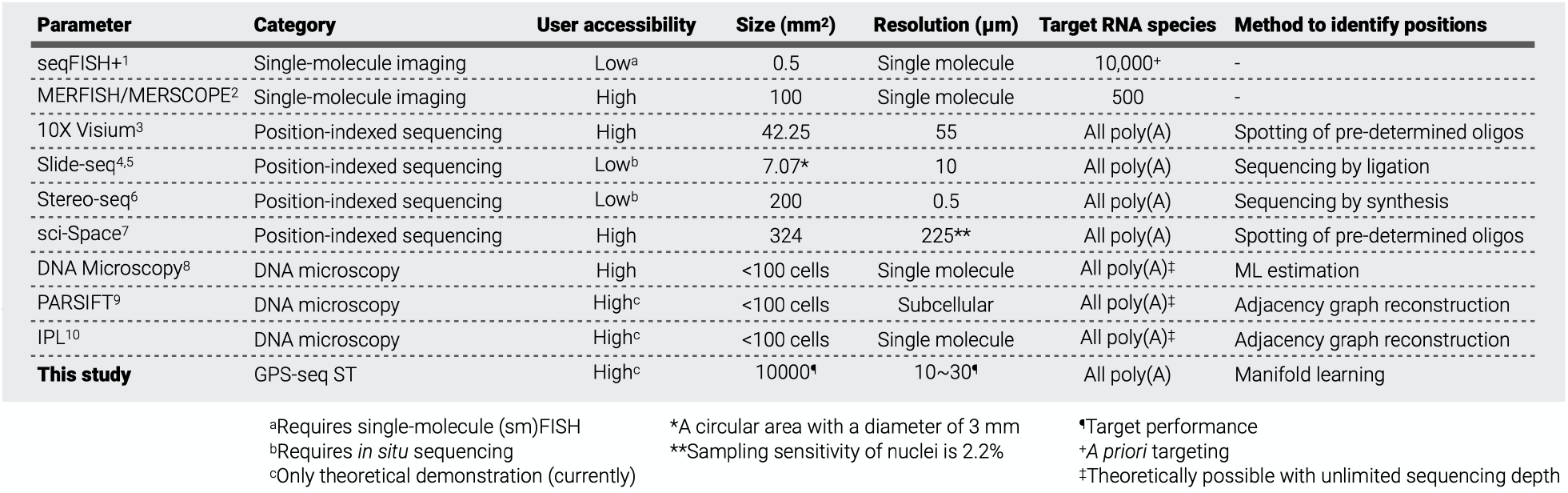
Comparison of spatial transcriptomics (ST) technologies.

Many existing ST technologies rely on optical microscopy, which suffers from an inherent trade-off between field-of-view and resolution. For example, fluorescence *in situ* hybridization (FISH) can resolve individual molecules of RNA through super-resolution imaging^15^. While FISH has classically been limited to resolving a handful of genes at a time, recent multiplex FISH strategies have expanded the number of targets^1,16^. However, in order to provide a precise quantitative estimate of RNA abundances, these methods count individual molecules within cells through super-resolution microscopy. This cannot be easily done over large spatial scales. Moreover, for FISH-based technologies, constructed probes cannot be utilized for other species. Spatial RNA sequencing has also been recently developed with the *in situ* sequencing by ligation (SBL) strategy^17,18^. Nevertheless, due to their reliance on optical microscopy, it is generally difficult to acquire large field-of-view with these single-molecule resolution methods. (While profiling of large-scale tissues is theoretically possible, it requires a time-consuming raster scan of a small field of view to cover the large area.)

Several technologies have recently achieved key steps towards large-scale spatial transcriptomics at single-cell resolution, based on ideas we refer to as “position-indexed sequencing”. This class of ST technologies employs spatially immobilized spots of DNA-barcoded reverse transcription (RT) primers to generate complementary DNA (cDNA) products conjugated with corresponding positional barcodes^3-7,19-21^. After deep sequencing, the spatial transcriptomic image can be reconstructed using the positional barcodes. However, the barcoded surfaces need to be prepared either by depositing a set of barcode primers of known sequences onto defined spots (e.g., 10X Genomics Visium^3^, sci-Space^7^ and DBiT-seq^19^) or by optical sequencing of barcode RT primers of unknown sequences after their dense surface deposition (e.g., Slide-seq^4,5,^, SeqScope^20^ and Pixel-seq^21^). The former suffers from spatial resolution and complexity in DNA spots, and the latter does not scale in spatial size due to the need for retrospectively identifying positional barcodes by *in situ* DNA sequencing using optical microscopy. If these barriers were overcome, it would revolutionize bioimaging and genomic understanding of animal bodies and their malfunctions (i.e., diseases).

Sequencing-based microscopy technologies have the potential to break this barrier because they do not rely on optical microscopic observation of specimens^8-10,22^. One approach, called DNA microscopy^8^, localizes RNA molecules through an *in situ* PCR reaction, where RNA molecules are reverse transcribed *in situ* with primers having unique molecular identifiers (UMIs) and PCR products of the UMI-tagged cDNA diffuse and make contact with other cDNAs. When contact is made, a unique event identifier is constructed by fusing UMI-tagged cDNA sequences from the different source RNA molecules. These event identifiers representing molecular proximity information are read out by high-throughput DNA sequencing, and the physical locations of all molecules are then reconstructed computationally from these data. In theory, this reaction can be performed at a tissue scale with a single readout by deep sequencing. However, this has only been demonstrated to date with images consisting of less than a hundred of cells. In practice, it would require tremendous sequencing depth and computing power to localize single molecules over large spatial scales.

While single-molecule resolution may eventually be useful, there is still much to be gained from studying tissue architecture at the level of cells, rather than individual molecules within cells. Indeed, some recent single-molecule studies have analyzed their data by aggregating them into the level of cells^1,6^. Here we propose a new “global positioning” approach to spatial transcriptomics, called GPS-seq, similar to the global positioning systems (GPS) used in navigation systems for cars and cell phones, where devices determine their precise location on Earth using their distance to multiple satellites (Fig. 1a). GPS-seq combines the advantages of sequencing-based microscopy and position-indexed sequencing. In GPS-seq, small satellite devices distribute unique nucleotide barcode molecules called satellite barcodes (sBCs) over a tissue which is applied on a surface of barcoded beads (Fig. 1b). These sBCs are collected by bead-specific RT primers which encode unique DNA barcodes, which we refer to as “anchor barcodes” (aBC) (Fig. 1c). The aBCs also collect transcriptome products of neighboring cells. After satellite and transcriptomic capture, the sample is subjected to cDNA synthesis that produces both aBC-sBC fused DNA products and aBC-transcript fused DNA products. After deep sequencing, the satellite barcode counts give sufficient information to localize the anchor barcodes, and therefore the associated transcriptome profiles. This strategy removes the current need to either deposit known positional barcodes at predefined locations or to determine positional barcodes with *in situ* sequencing. This enables an unprecedented scale of spatial transcriptomics at single cell resolution.

**Figure 1.**
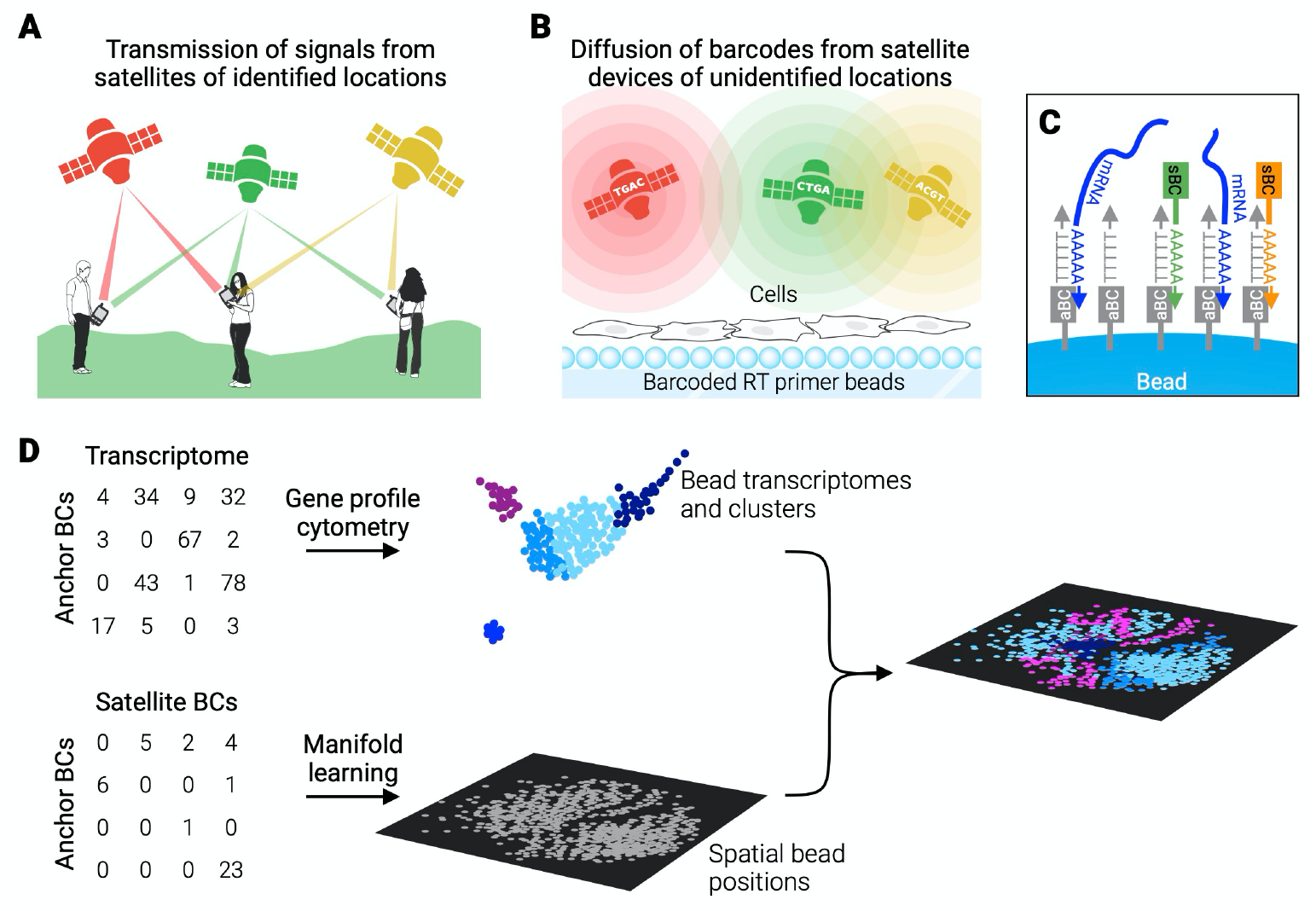
GPS-seq. (**A**) The Global Positioning System (GPS). (**B**) GPS-seq for scalable high-resolution spatial transcriptomics (ST). GPS-seq reconstructs spatial positions of barcoded beads of unknown positions using only deep sequencing and manifold learning. (**C**) Small beads coated with reverse transcription (RT) primers with a unique anchor barcode (aBC) capture both neighboring mRNAs and polyadenylated satellite barcode (sBC) nucleotides. The sBC and transcriptome sequences are fused to the aBCs by RT for deep sequencing. (**D**) Large-scale ST reconstruction from only sequencing data. Spatial positions of barcoded beads are reconstructed from their sBC abundance profiles using manifold learning. Transcriptome profiles of barcoded beads are then mapped to the spatial positions.

The principles of GPS-seq are quite general and could be realized by different experimental strategies for transmitting and capturing satellite barcodes and genomic information. For example, instead of capturing satellite barcodes on a two-dimensional surface of beads, one could transmit satellite barcode molecules directly onto cells in a tissue and then profile either gene expression or chromatin accessibility via standard droplet-based or split-pool scRNA-seq or scATAC-seq. In such cases, the single-cell barcodes can serve as “anchor barcodes”. For concreteness, we refer to the beads approach throughout most of the manuscript (e.g., we refer to “localizing beads” and discuss how many “reads per bead” are necessary to achieve single-cell localization accuracy). In the remainder of this manuscript, we outline the theoretical principles and scalability of GPS-seq. We determine the optimal physical properties of satellite barcode transmission systems and quantify the resolution and field of view that can theoretically be achieved by sequencing satellite barcode libraries at different depths.

## Results

### GPS-seq localizes beads from satellite barcodes via manifold learning

In ordinary GPS, a cell phone determines its position from distances to multiple satellites, whose positions are known in advance. Even if the locations of the satellites were unknown, you could still determine your position relative to the satellites, as well as relative to someone else who received signals from some of the same satellites. We leverage these ideas in GPS-seq to determine the positions of barcoded beads, without *a priori* establishing the positions of the devices which distribute satellite barcodes. The GPS-seq measurement process produces two data matrices—one containing counts of sBCs for each aBC, and another containing counts of transcription products for each aBC (Fig. 1d). We use the matrix of sBC counts to determine the positions of the beads, and then fill in the data from the second matrix to form the spatial-transcriptomic image. Here we focus on this first step of determining the physical positions of the beads from the sBC data matrix. For a detailed mathematical model of the measurement process, see Supplementary Equations.

The core principle of GPS-seq is that beads which are nearby in physical space will collect similar counts of satellite barcode molecules because satellite barcodes are distributed in a spatially coherent way—they diffuse locally from satellite devices. In mathematical terms, the measurement process can be understood to embed the physical two-dimensional surface of beads into a high-dimensional “satellite barcode space,” where each bead is represented by its vector of sBC counts, similar to how a transcription profile is represented by a high-dimensional gene expression vector in the analysis of scRNA-seq data. Our key insight is that the embedded beads form a two-dimensional manifold, or surface, in high-dimensional sBC space (Supplementary Equations, Extended Data Fig. 1). Thus, our goal is to reverse the embedding and recover the original positions of the beads by “learning the manifold”. Manifold learning is a well-established field^23-26^, and is routinely applied to visualize high-dimensional single-cell RNA-seq data^27^. While the interpretation of these visualizations is usually related to an abstract “manifold of cell states”, we have here a very concrete and physically interpretable setup: the two-dimensional surface on which beads are deposited (Fig. 2a) is embedded into the high-dimensional sBC count space (Fig. 2b). Thus, in the following, we tested whether manifold learning algorithms could recover the physical positions of beads from the sBC data (Fig. 2c) in various conditions.

**Figure 2.**
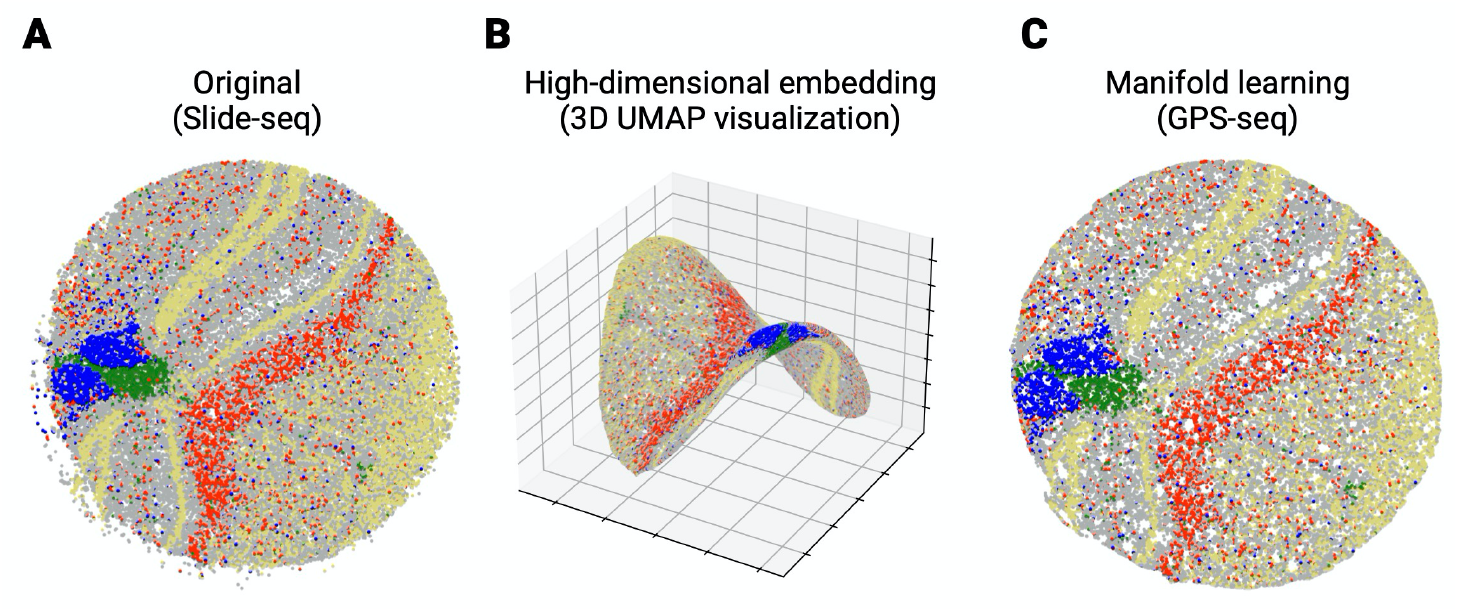
Manifold learning of spatial bead positions. (**A**) Original Slide-seq hippocampus data colored by cell type. (**B**) Illustration of beads in a high-dimensional satellite barcode space. In GPS-seq, bead positions are embedded in a many thousand-dimensional sBC space, where the surface of beads forms a two-dimensional manifold. Here we visualize this two-dimensional manifold in three-dimensional space via UMAP. (**C**) The Slide-seq data can be reconstructed using GPS-seq, without using the bead position information using manifold learning. Reconstruction parameters: 100,000 satellites/cm^2^, 50 μm diffusion, 130 UMIs per bead (556 RPB).

### *In silico* demonstration

We performed extensive simulations in order to determine the optimal properties of GPS-seq satellite devices. *How many satellite devices should we have? What level of diffusion is optimal for the satellite barcodes? How many reads per bead do we need to sequence?* We found that we could achieve single-cell resolution over a wide range of parameters. We simulated randomly positioned satellite devices producing unique sBCs over bead positions from Slide-seq datasets^5^ as well as synthetic datasets with hundreds of thousands of beads (Methods). We modeled satellite devices as producing a Gaussian profile of sBCs where the width of the Gaussian could be varied to simulate varying levels of diffusion (Fig. 3a). Each bead captures reads from nearby sBCs, proportional to the intensity of the sBC diffusion profile at the bead’s position.

**Figure 3.**
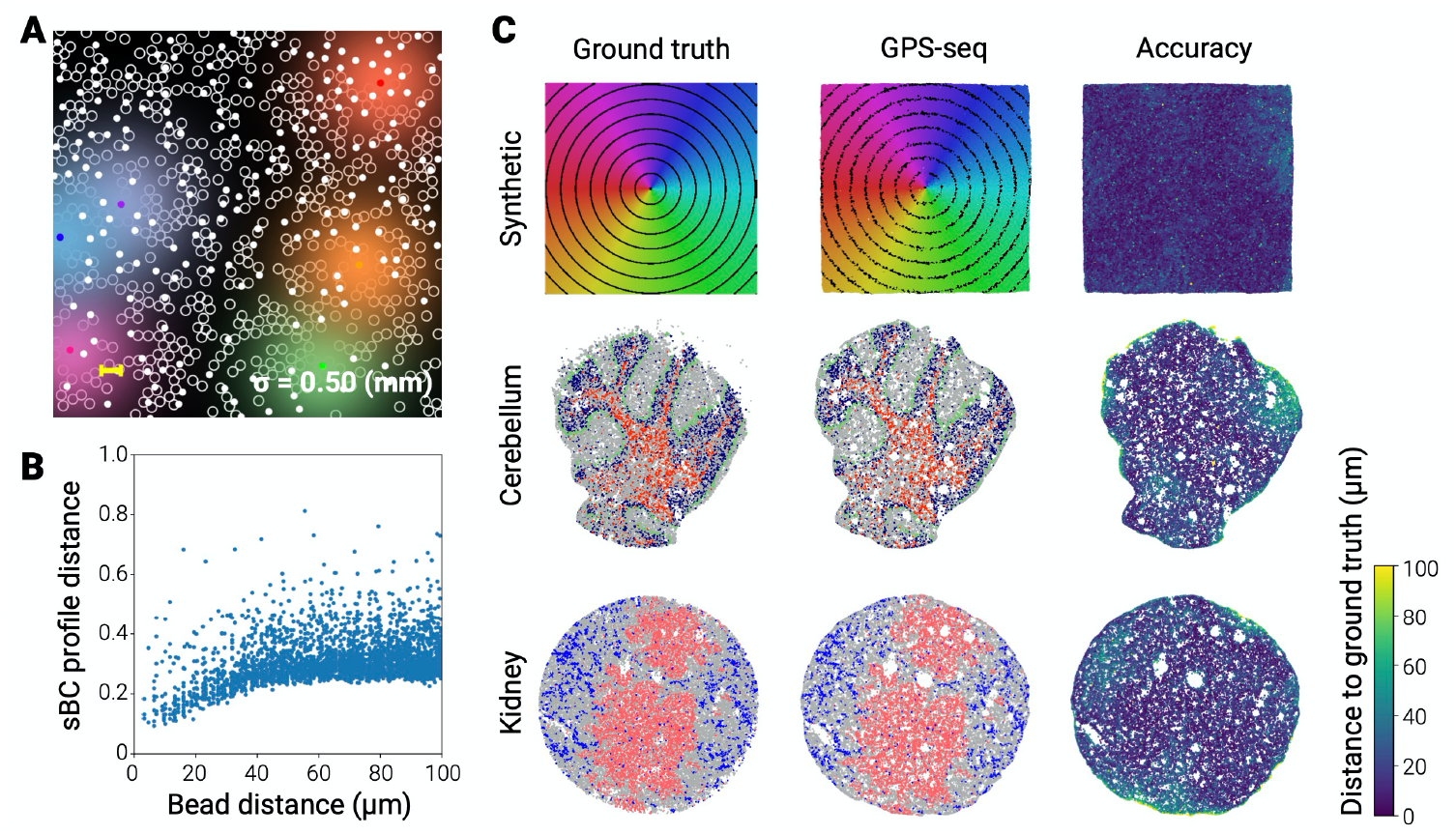
Single-cell resolution with manifold learning. (**A**) An illustration of satellite devices distributed over barcoded beads (white open circles) based on densities in the Slide-seq simulations. The sBC diffusions (blurred colors) are represented for some of the arbitrary picked satellites. (**B**) Correlation of physical distances and sBC profile distances for 10^5^ randomly selected pairs of beads from the synthetic dataset. (**C**) GPS-seq reconstructions (center, right) compared to the ground truth (left) for the synthetic dataset and two Slide-seq datasets. All reconstructions achieve a median reconstruction distance below 20μm using 100,000 satellites/cm^2^, 50 μm diffusion and 130 UMIs per bead (556 reads per bead).

In practice, beads can vary in quality, with higher quality beads capturing more reads and *vice versa*. We modeled this data bias by down-sampling of reads from the Slide-seq kidney dataset (Methods, Extended Data Fig. 2a). Furthermore, groups of satellites may share the same barcodes, due to finite and biased sBC complexity, and reads from satellites sharing the same barcode are indistinguishable. We accounted for this in our simulations by randomly selecting a barcode for each satellite device from a pool of possible barcodes according to an experimental distribution of barcode frequencies (Methods, Extended Data Fig. 2c). This corresponds mathematically to a random projection in sBC space (Supplementary Equations). However, random projections are known to preserve pairwise distances in high-dimensions^28^. Indeed, when we examined the Euclidean distance between beads in “projected” sBC count space, we found that beads that are spatially close together share similar sBC vectors and were close together in high-dimensional space (Fig. 3b). This correlation understandably dropped off at large-scale distances, because sBCs only diffuse to a limited extent and any pair of beads that share no satellites in common are roughly the same distance apart in the sBC count space.

We found that the UMAP algorithm^25^ can leverage local distances in sBC count space to reconstruct the positions of aBC beads at single-cell resolution. Using the data biases in read count per bead (rpb) and sBC redundancy described above, we tested GPS-seq on a synthetic dataset and two Slide-seq datasets to get an idea for how reconstructions look on real tissue samples (Fig. 3c). We swept over both UMAP hyperparameters and physical parameters (number of satellite devices and sBC diffusion levels) at multiple levels of median rpb (Methods). We aligned the output of UMAP to the ground truth and computed the distance of each bead to its true position. In Figure 3c, we show results for parameter settings where GPS-seq performs well at low sequencing depth: we can achieve a median distance to the ground truth below 20 μm using 556 rpb (or 130 UMIs per bead) and satellite diffusion coefficient σ=50 μm in all three cases.

### Robustness and scalability of GPS-seq

Further, we found the method can achieve single-cell resolution across a wide range of physical parameters. We display the results of this sweep on synthetic data in Figure 4. Our best reconstructions achieved a median distance per bead to the ground truth of 10 μm, requiring only 180 UMIs per bead (1,110 rpb). Sequencing depth can be significantly reduced by moderately decreasing the resolution, with our method achieving reconstructions under 30 μm with as few as 15 UMIs per bead (30 rpb). As the size of a mammalian cell varies from 10-100 μm^29,30^, both reconstructions are near the low end of the single-cell range. Further, our method is robust to changes in physical parameters (Extended Data Figs. 3-7). Testing diffusion levels from 30-100 μm, numbers of satellite devices from 25,000-250,000 per cm^2^, and varying the sequencing depth up to 265 UMIs per bead, we achieved median alignment errors under 20 μm for all but one combination of diffusion level and number of satellites (Extended Data Fig. 3). Specifically, we achieve 10 μm resolution reconstructions with sBC diffusion from 30–50 μm and 100,000–250,000 satellite devices per cm^2^. We achieved 30 μm resolution over an even wider range, with sBC diffusion from 30–100 μm and 25,000–250,000 satellites per cm^2^.

**Figure 4.**
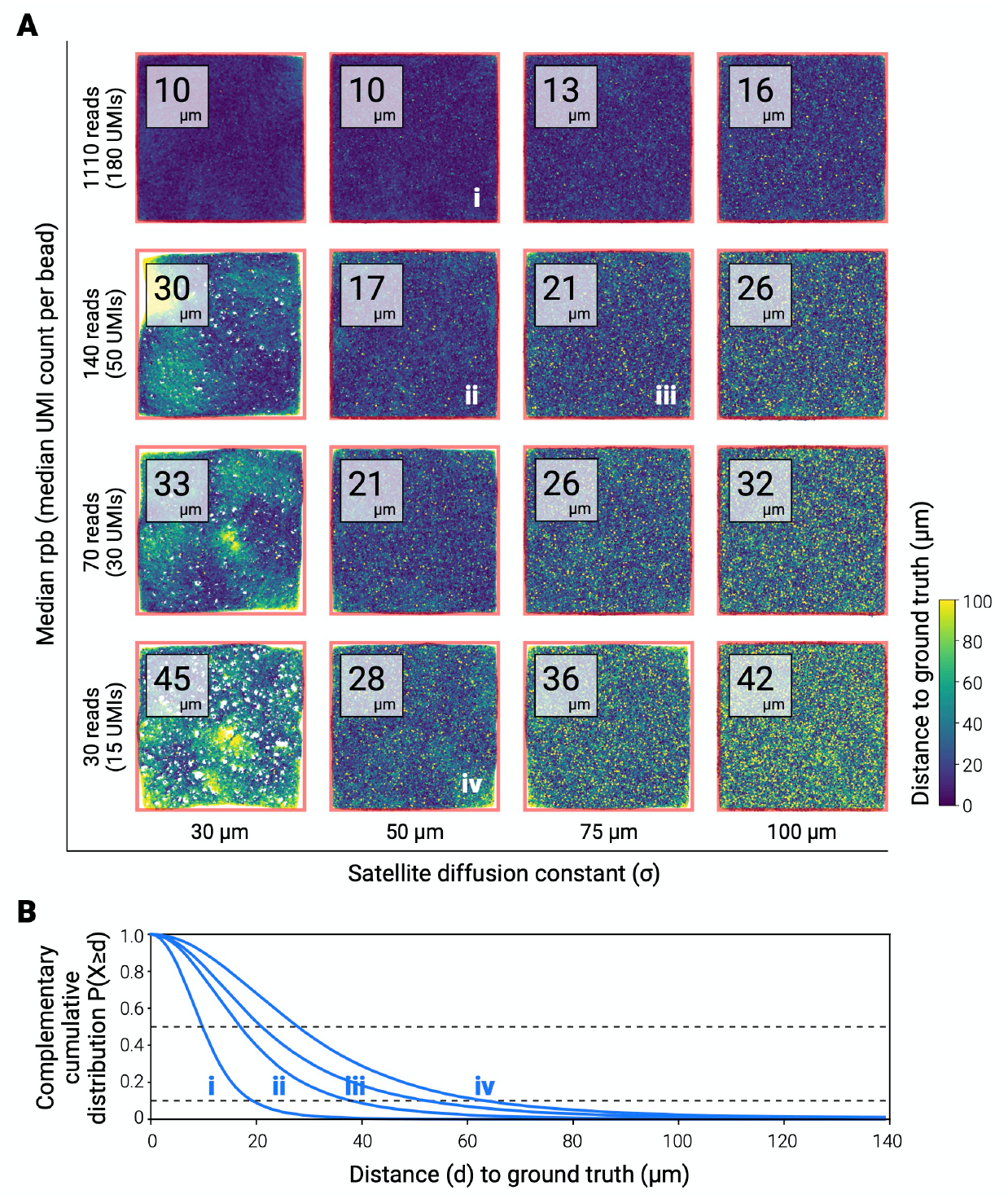
Single-cell resolution is possible over a range of physical parameters. (**A**) The best reconstruction across satellite densities and UMAP parameters on the synthetic dataset for each combination of sequencing depth and diffusion level. Images show the distance to the ground truth for each bead with the median distance inset. (**B**) Complement of the cumulative distribution of bead distances to ground truth for each of the four reconstructions indicated in (A). Dashed lines indicate the 50th and 90th percentiles of beads.

Finally, we sought to compare the resolution and scalability of GPS-seq to other state-of-the-art ST technologies. As we sequence sBCs more deeply, we are able to localize beads more precisely (Fig. 5a). Using this plot, we estimated the resolution (μm) and field-of-view (image width of a square in mm) achieved by GPS-seq for different total sequencing depths and compared them to seven leading methods (Fig. 5b). For GPS-seq, we show contours achievable for several different total sequencing depths, as the resolution of GPS-seq depends on the depth of sequencing. Intuitively, the number of reads per bead varies along each curve. With 10 billion reads (one NovaSeq run), GPS-seq can localize 10^8^ beads to generate a 10 cm by 10 cm image with single-cell resolution (17 μm).

**Figure 5.**
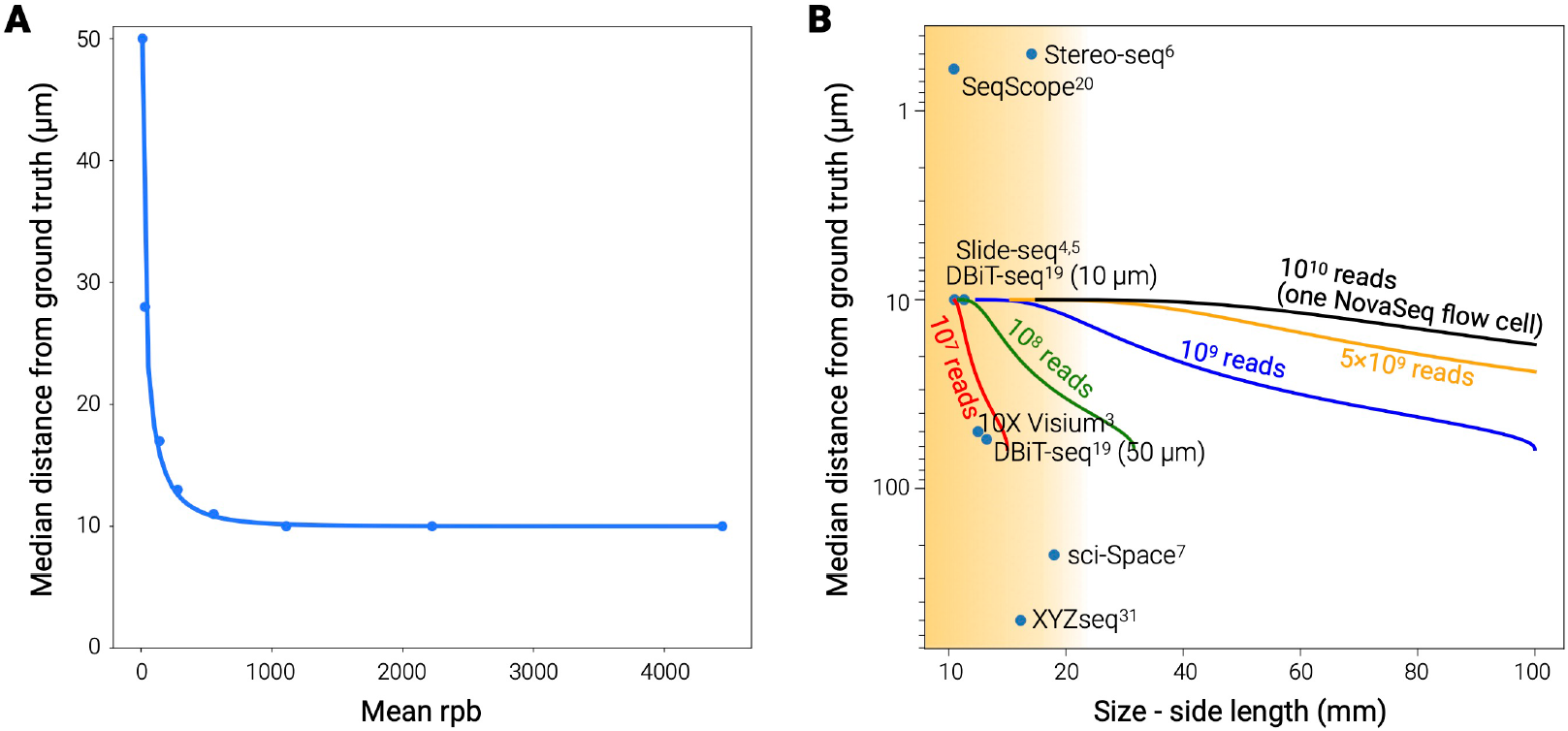
Comparison to existing ST technologies. (**A**) Best median reconstruction resolution versus mean reads per bead obtained from the simulated data sweep with an exponential line of best fit. (**B**) Performance of GPS-seq versus current ST technologies. Lines indicate interpolated GPS-seq performance using the curve fit in (A) and assuming densely packed 10 μm beads. Labeled dots indicate existing ST technologies, with the shaded region indicating the current achievable scale.

## Discussion

We envision that GPS-seq can be practically implemented with different strategies for satellite barcode systems (Fig. 6). The three general components that we require are:

**Figure 6.**
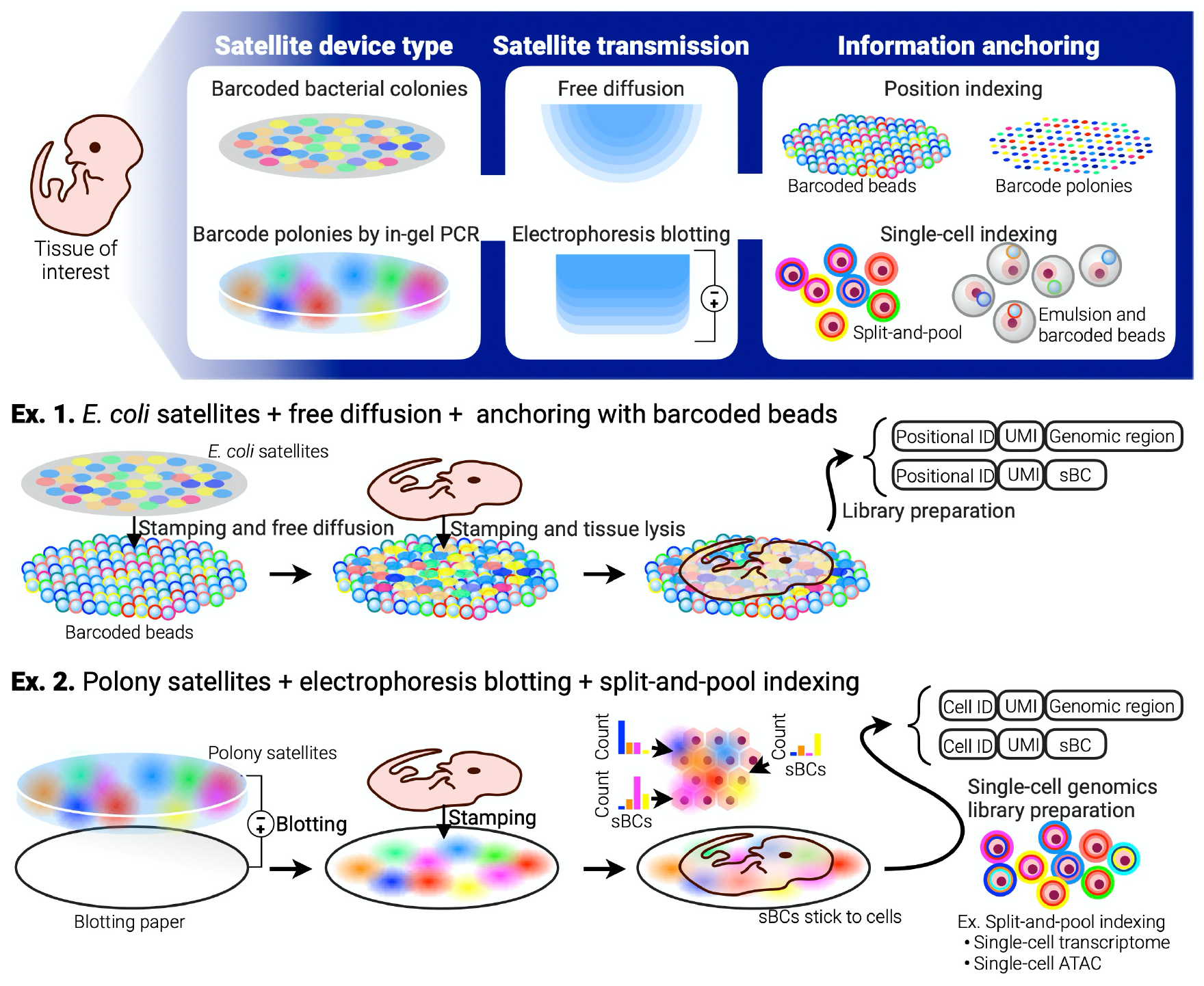
Insights on implementing GPS-seq satellite barcode systems. The potential implementations of GPS-seq can be categorized into three modules: (1) satellite device type, (2) satellite transmission method and (3) information anchoring method. We envision that satellite devices could be implemented by either engineering bacterial colonies each expressing unique sBCs or seeding a gel with unique sBC DNA molecules and performing in-gel PCR to amplify the sBCs to produce polymerase colonies (polonies). Satellite transmission could be achieved by either free diffusion or electrophoresis blotting. Finally, anchor barcodes could be implemented by either capturing RNA on positional barcodes such as barcoded beads (or on barcoded spots) or with standard single-cell genomics methods. **Ex. 1:** *E. coli* colonies serve as satellite barcodes and a surface of barcoded beads serves as anchor barcodes. The colonies and then the tissue sample are stamped onto the beads. **Ex 2:** diffused satellite barcodes can be directly deposited to a target tissue section, where single-stranded sBC molecules stick to cells (and nuclei). After dissociation of the satellite-attached tissue into single cells, any of the single-cell genomics platforms can conjugate single-cell RNAs (or DNAs) and their sBC profiles to the single-cell identifiers, enabling scalable spatial transcriptomics with single-cell resolution.

1. Satellite devices produce unique sets of satellite barcode molecules. The numbers of satellite devices should be comparable to the number of cells. Some barcode overlap can be tolerated.
2. Satellite barcodes are transmitted to the tissue sample. Some diffusion is beneficial so that beads receive sBCs from multiple neighboring satellites. The optimal diffusion level may be ∼50 μm for 10 μm beads.
3. sBCs are captured together with genetic material from cells, and both are fused to aBCs (i.e., positional barcodes or single-cell barcodes). This “information anchoring” allows a ST image to be formed after sBCs are used to localize aBC compartments.

Large arrays of satellite devices could be prepared by either:

a. engineering bacterial cells to express unique sBCs and seeding them sparsely to obtain clonal sBC-expressing colonies of up to 50-μm diameter (Fig. S8),
b. seeding a gel with unique sBC DNA molecules and performing in-gel PCR to amplify the sBCs to produce “polymerase colonies”^32^.

These satellite devices could be transmitted to the tissue by either free diffusion or electrophoresis blotting, and the diffusion level of sBCs could be tuned during this step. Finally, anchor barcodes could be implemented by capturing RNA on positional barcodes such as barcoded beads (or on barcoded spots) as we outline in Ex. 1 of Figure 6. However, the general principles of GPS-seq are also compatible with standard single-cell genomics methods such as droplet microfluidics or split-pool indexing. As illustrated in Ex. 2 of Figure 6, diffused satellite barcodes can be directly deposited to a target tissue section, where single-stranded sBC molecules stick to cells (and nuclei) ^7^. After dissociation of the satellite attached tissue into single cells, any of the single-cell genomics platforms can conjugate single-cell RNAs (or DNAs) and their sBC profiles to the single-cell identifiers, enabling scalable spatial transcriptomics with single-cell resolution. Theoretically, any modality of high-coverage single-cell genomics, such as sci-ATAC-seq, could be transformed into spatial genomics in this strategy as long as sBCs can be captured by single-cell identifiers together with genomic materials of interest.

Accordingly, GPS-seq makes it possible to profile large-scale tissues at single-cell resolution, orders of magnitude larger than is possible with current technologies. With this technology, neuroscientists could chart the spatial transcriptomic architecture of entire brains. Oncologists could profile spatial patterns of gene expression across entire tumors, revealing the spatial structure of the tumor microenvironment. In drug discovery, one could profile a cross-section of a whole adult mouse to systematically characterize the effects of a compound across the body. Tissue engineers could gain new insights into the spatial composition of synthetic tissues and organs. High-throughput deep sequencing and single-cell genomics technologies have rapidly advanced and triggered paradigm shifts in biology in the last decades. While none of the current ST technologies are currently scalable and largely accessible by the community, we foresee that our new theoretical framework will overcome the current technical difficulties and lead the field like the previous genomics technologies.

## Acknowledgements

We thank members of the Schiebinger lab and Yachie lab for valuable discussions and critical assessment of the work, especially Rebecca Bonham-Carter for the help with the theorem proof. This study was supported by the Canada Research Chair program (by the Canadian Institutes for Health Research) to N.Y., the Career Award at the Scientific Interface from the Burroughs Wellcome Fund to G.S., and the Genome British Columbia’s Pilot Innovation Fund program (PIF003) to N.Y. and G.S. Y.K. was supported by the Japan Society for the Promotion of Science (JSPS) Research Fellowships.

## Author contributions

N.Y. and Y.K. conceived the high-level concept of GPS-seq. G.S., L.G., A.A., Y.K., and M.H. conceived the use of manifold learning. N.Y. and G.S. designed the study. L.G., A.A., Y.K. and M.H. led the analyses. S.I. and S.K. supported the theoretical data analyses. L.G., A.A., Y.K., N.Y., and G.S. wrote the manuscript.

## Data and Code Availability

All the codes used in this study are available at https://github.com/schiebingerlab/GPS-seq

## Methods

### Simulated data generation

To generate data, we chose ground truth positions for beads and randomly generated satellite positions extending slightly beyond the beads to ensure uniform reads. We used ground truth positions from the Slide-seq kidney, hippocampus, and cerebellum 2 datasets as well as dense grids of 100,000 beads to test scalability. We modelled the reads from each satellite as following a Gaussian distribution where the width of the Gaussian determined the level of diffusion. In addition to diffusion, the number of satellites was a hyperparameter for data generation.

### Bead quality and barcode overlap

To sample an sBC read matrix we needed to define the distributions of satellite barcode redundancy and bead quality (see Supplementary Equations for mathematical details). To account for varying bead qualities, we utilized the distribution present in the Slide-seq kidney dataset (See Extended Data Fig. 2a). Slide-seq datasets were downloaded from Single Cell Portal. cDNA sequences were extracted from the BAM file and stored as FASTQ format using samtools [33]. The reads were downsampled using seqkit [34] and mapped to the reference genome version of mm10 to obtain a UMI count matrix for each downsampling level. This resulted in the mean rpb to UMI curves shown in Extended Data Fig. 2b. We found that the read to UMI curve differed between different Slide-seq samples. As our model depends on UMI while sequencing cost depends on reads, we provide both values in our results.

Barcodes do not occur uniformly in practice leading to some satellites sharing common barcodes. We quantified the extent of this phenomenon by conducting the following experiment. We used a barcoded plasmid library that has been previously generated [35]. To generate the high-throughput sequencing library, the plasmid pUC119 encoding eight randomized nucleotides (4^8^ possible barcode combinations) was subjected to PCR amplification using Phusion DNA polymerase (NEB). After purification of the first PCR product, we reamplified the sample with an index primer using Phusion DNA polymerase (NEB). The sequencing sample was quantified using a KAPA Library Quantification Kit Illumina (KAPA Bioscience) and sequenced by Illumina HiSeq2000 with 20% PhiX spike-in control (Illumina). The sequencing reads were demultiplexed according to the sample indices and constant sequences using NCBI Blast+ (version 2.6.0) with the blast-short option. Using read alignment information, 8-mer barcodes were extracted to count each barcode abundance. The resulting barcode occurrence distribution is shown in Extended Data Fig. 2c. We sampled satellite barcodes from this distribution for our simulations.

### Forming the sBC Count Matrix

To create the sBC count matrix we first form a matrix with no barcode redundancy where for each bead we sample a bead quality from the Slide-seq kidney distribution and then sample the specified number of reads from the sBC distribution determined by the bead’s position (See Supplementary Equations for mathematical details). We then generate a barcode for each satellite, following the experimental distribution, and combine columns of the initial matrix sharing a barcode. Our data generation process has three hyperparameters: (1) number of satellites, (2) diffusion level, and (3) sequencing depth. We tested a grid of four numbers of satellites, # satellites ∈ [25000, 50000, 100000, 250000]*/*cm^2^, six diffusion levels, *σ* ∈ [10, 20, 25, 30, 50, 75, 100] *µm*, and 11 downsampling levels, mean rpb ∈ [10, 30, 50, 70, 90, 140, 280, 555, 1100, 2225, 4445].

### Reconstructions with Manifold Learning

UMAP is a dimensionality reduction technique that is frequently used with single-cell data to reduce a gene expression matrix to two dimensions for visualization. When applied to expression data, the embedding captures structure in gene expression space, often grouping cells with similar biological function or at similar stages of development. However, by applying it to our sBC matrix we can recover the underlying two dimensional manifold.

UMAP works by first building a graph representation of the high-dimensional data, using local metrics around each data point. It then optimizes a graph in the low-dimensional space so that it is structurally close to its high-dimensional counterpart, by minimizing the cross-entropy between the reconstructions. Working with local distances is critical for our application, as with limited sBC diffusion and read depth, distances are only meaningful between beads that have reads from common satellites. Working with local distances also facilitates scalability, as the distance between most pairs of beads is not considered, preventing the algorithm from scaling quadratically. As UMAP allows an arbitrary distance function, we tested Euclidean distance on raw counts, row-normalized, and log-tpm normalized data as well as cosine similarity and found the row-normalized Euclidean distance resulted in the best reconstructions.

UMAP introduces two hyperparameters: (1) the number of neighbors to consider for each bead, *n_neighbors*, and (2) the minimum distance between points in the low-dimensional space, *min_dist*. We tested a grid of four values for each hyperparameter, *min_dist* ∈ [0.25, 0.5, 0.75, 1] and *n_neighbors* ∈ [25, 50, 75, 100], for every combination of hyperparameters from our data simulation. The reconstruction was not highly sensitive to the number of neighbors with *<* 15*µm* alignment distances being possible from 10-60 neighbors, with the Slide-seq data performing best around 20 neighbors and the dense grids of beads performing best around 60 neighbors. As higher values of *min_dist* tend to result in more uniform embeddings, we found values of 0.75 and 1 tended to perform best on our data, especially for our best embeddings. However, in some cases we did find lower values of *min_dist* resulted in the best reconstruction.

As we used simulated data, we were able to quantify our performance relative to the ground truth, which would not be possible with experimental data. However as we expect a nearly uniform distribution of beads, the uniformity of the embedding could be used to evaluate the performance of UMAP hyperparameters in practice.

### Reconstruction Alignment

As distances passed to UMAP are relative, the embedding may be a rotation, reflection, scaling, or translation of the ground truth. We aligned each embedding to the ground truth using the Kabsch-Umeyama algorithm [36], which finds the rigid transformation (rotation, reflection, scaling, and translation) that minimizes the sum of distances between corresponding points in the embedding and the ground truth.

### Formation of bacterial satellite colonies

We have recently established the protocol to form tiny, dense satellite bacterial colonies on a paper disc of any size. In this protocol, a paper disc is soaked into an E. coli cell culture and placed on a solid LB (Luria Broth) agar plate where small bacterial colonies are formed densely but with minimal between-colony contacts. We performed the experiment using fluorescent protein-expressing cells (Extended Data Figure 8) and demonstrated that this protocol can confer small sizes of dense E. coli colonies whose diameters are up to 40 µm.

### Generation of complex DNA barcode library

We have also established the protocol to generate a complex barcoded E. coli library. Briefly, a DNA oligo pool having a random sequence region is amplified by PCR and ligated into a backbone vector. The ligation product is transformed into chemically competent cells and densely plated on solid LB agar plates, allowing cell growth. After applying liquid LB media or Tris-EDTA, the bacterial lawn is scraped and collected. PCR amplification and deep sequencing of the DNA barcodes from the cell pool enables the assessment of library complexity (Extended Data Figure 2c). We previously achieved a barcode complexity of 6.7810^5^.

### Plasmid construction and generation of fluorescent protein expressing cells

We cloned a gene cassette encoding an EGFP, sBC cloning site and poly(A) sequence together with the J23110 promoter sequence into the pUC-19 backbone plasmid by Gibson assembly. For the red fluorescent protein expressing plasmid, we replaced EGFP in the EGFP expressing vector with mCyRFP1. Both assembled constructs were respectively transformed into NEB5a E.Coli competent cells, and the overnight-incubated colonies on solid LB+Agar plates were isolated followed by sequence integrity validation with Sanger sequencing.

### Satellite colony formation

To densely form bacterial satellite colonies on a two-dimensional surface, we utilized a paper disc (a blue separation paper for membrane filters: Millipore Sigma GTTP02500) as a scaffold of colony growth. We mixed an equal amount of EGFP and mCyRFP1 expressing cells cultured in LB+Ampicillin and soaked the paper disc into the mixture. After wiping out the excess liquid from the disc, we placed it on a solid LB agar plate and incubated it overnight at 37°C.

### Observation of satellite colonies

We observed the satellite colonies covered on the paper disc using Keyence BZX710. Images were processed by FIJI ImageJ Version 2.1.0/1.53c to quantify the numbers and areas of satellite colonies.

## Extended Data

**Extended Data Fig. 1:**
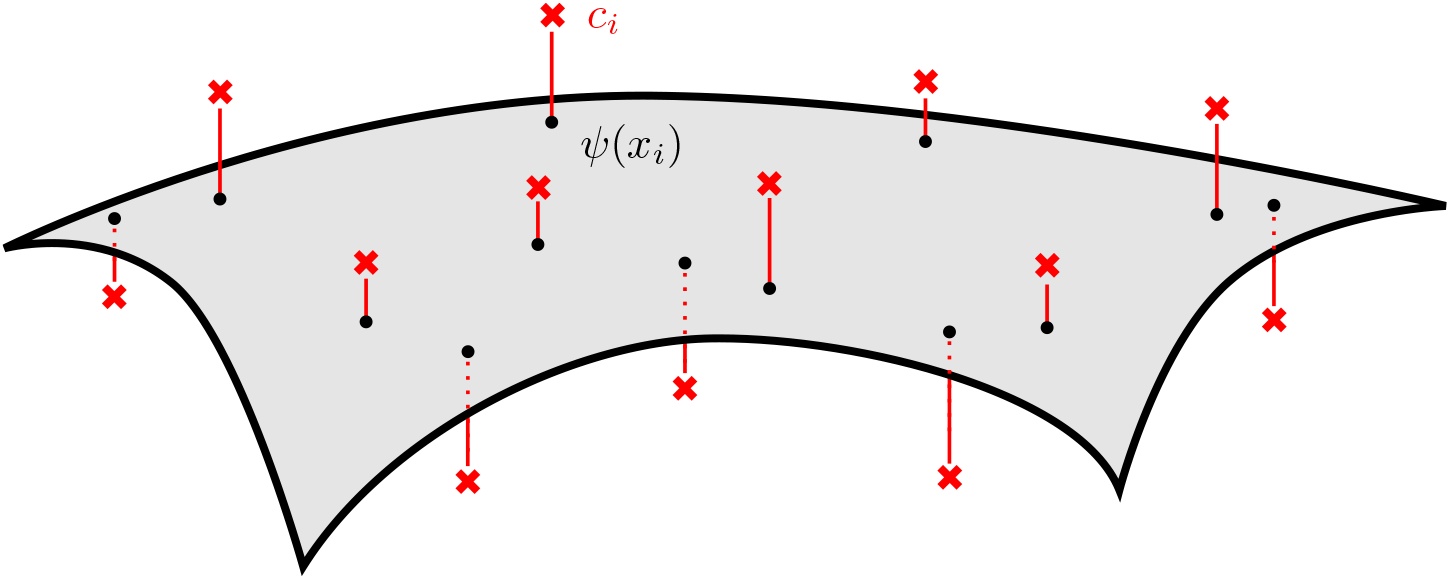
The two-dimensional manifold of beads, ℳ, embedded in satellite barcode space 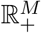. While the positions of the beads in satellite barcode space, *ψ*(*x*_*i*_), belong to ℳ, due to perturbations introduced by sampling and bead quality the observed counts, *c*_*i*_, are displaced from ℳ by a small amount of additive noise.

**Extended Data Fig. 2:**
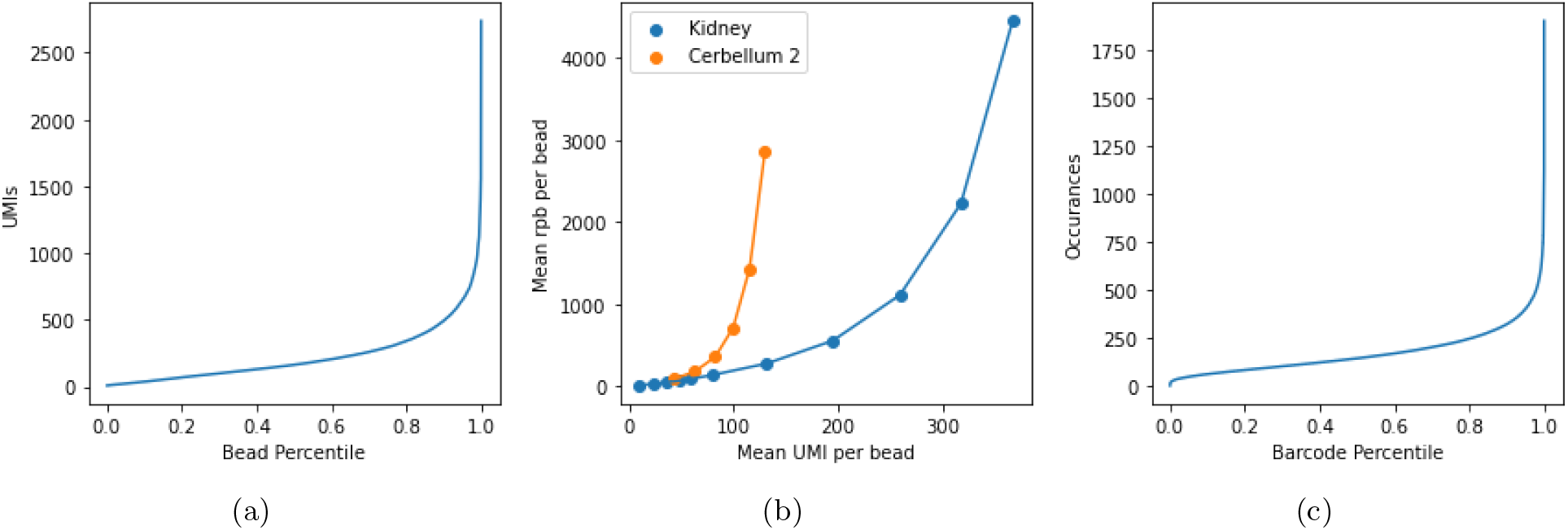
(a) Bead quality distribution for the Slide-seq kidney dataset. (b) Mean UMI versus rpb on downsampled versions of the Slide-seq kidney and cerebellum 2 datasets. While the curves are similar for low UMI, the diverge at high UMI values. As our method depends on UMI but sequencing cost depends on the number of reads, we report both UMI and the number of reads required in the Kidney dataset throughout the text. (c) Experimental barcode redundancy distribution.

**Extended Data Fig. 3:**
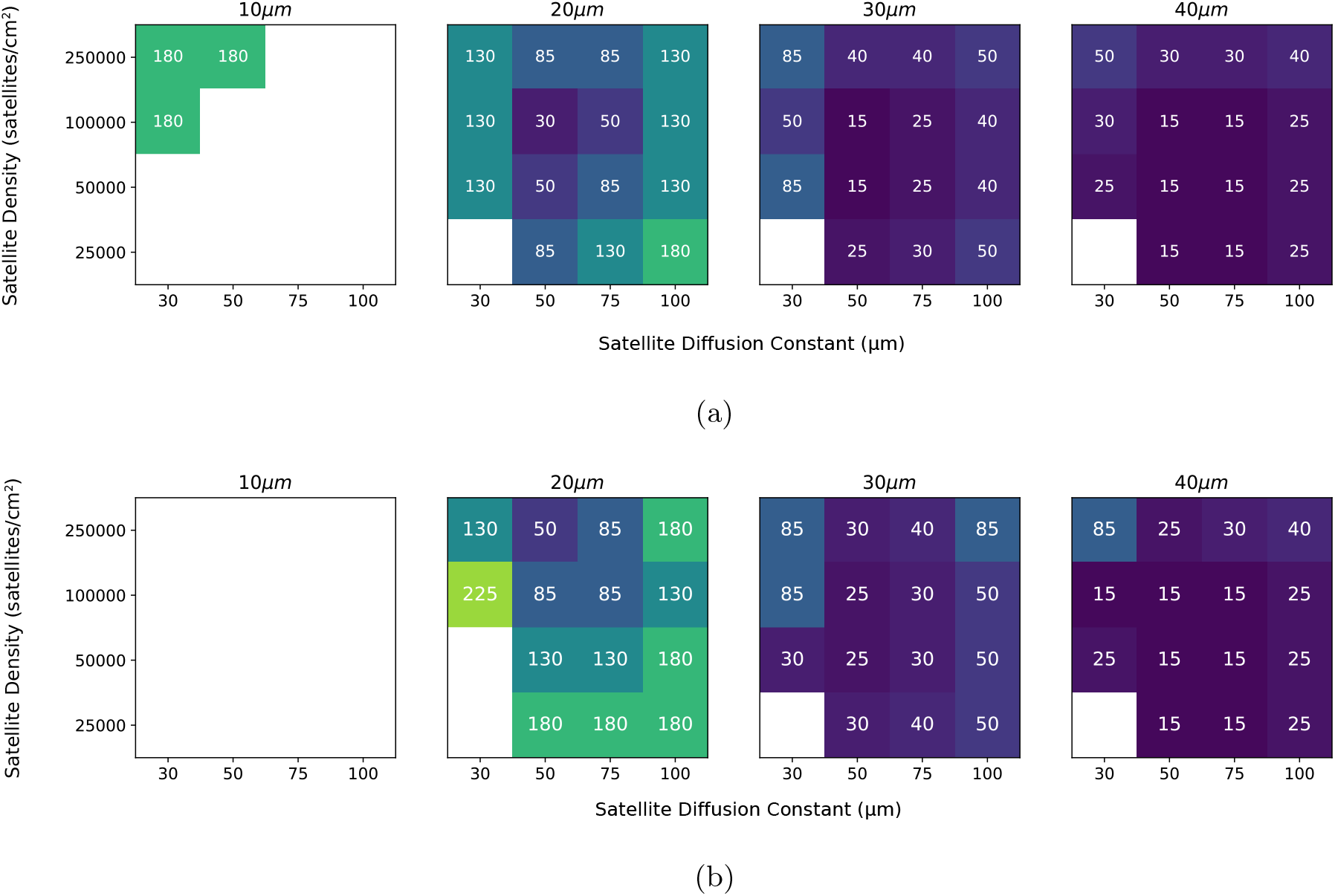
The value in each cell shows the minimal median UMI needed to reach a specific localization error (i.e. median alignment error, which is indicated above the block in *µm*). Each block shows how required sequencing depth varies with number of satellites and diffusion level for reconstructions on (a) 100,000 beads on a regular grid and (b) bead positions from the Slide-seq kidney dataset. Empty cells indicate no UMI up to 265 achieved that reconstruction quality for that combination of physical parameters. These reconstructions show that we can achieve at the low end of the single cell range (10 - 100 *µm*) for a wide range of physical parameters and that the sequencing depth can be varied to reach a target resolution for the lowest sequencing cost.

**Extended Data Fig. 4:**
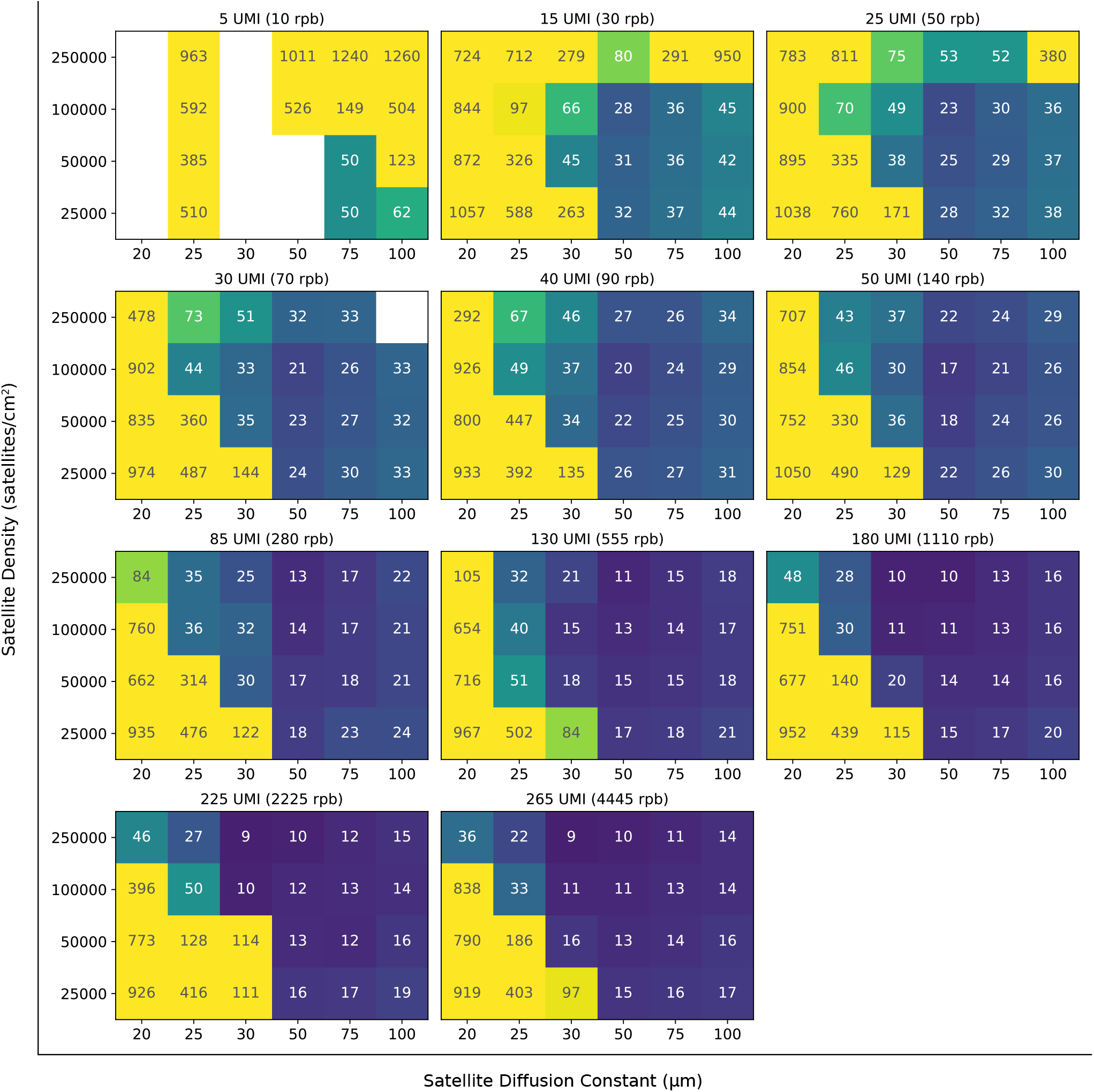
Summary of median alignment values in *µm* by number of satellites, diffusion level, and sequencing depth for simulations of 100,000 beads on a regular grid. We achieve at the low end of single cell resolution (10 - 100 *µm*) for a wide range of physical parameters. Further, sequencing depth can be adjusted to achieve a target resolution at the lowest sequencing cost.

**Extended Data Fig. 5:**
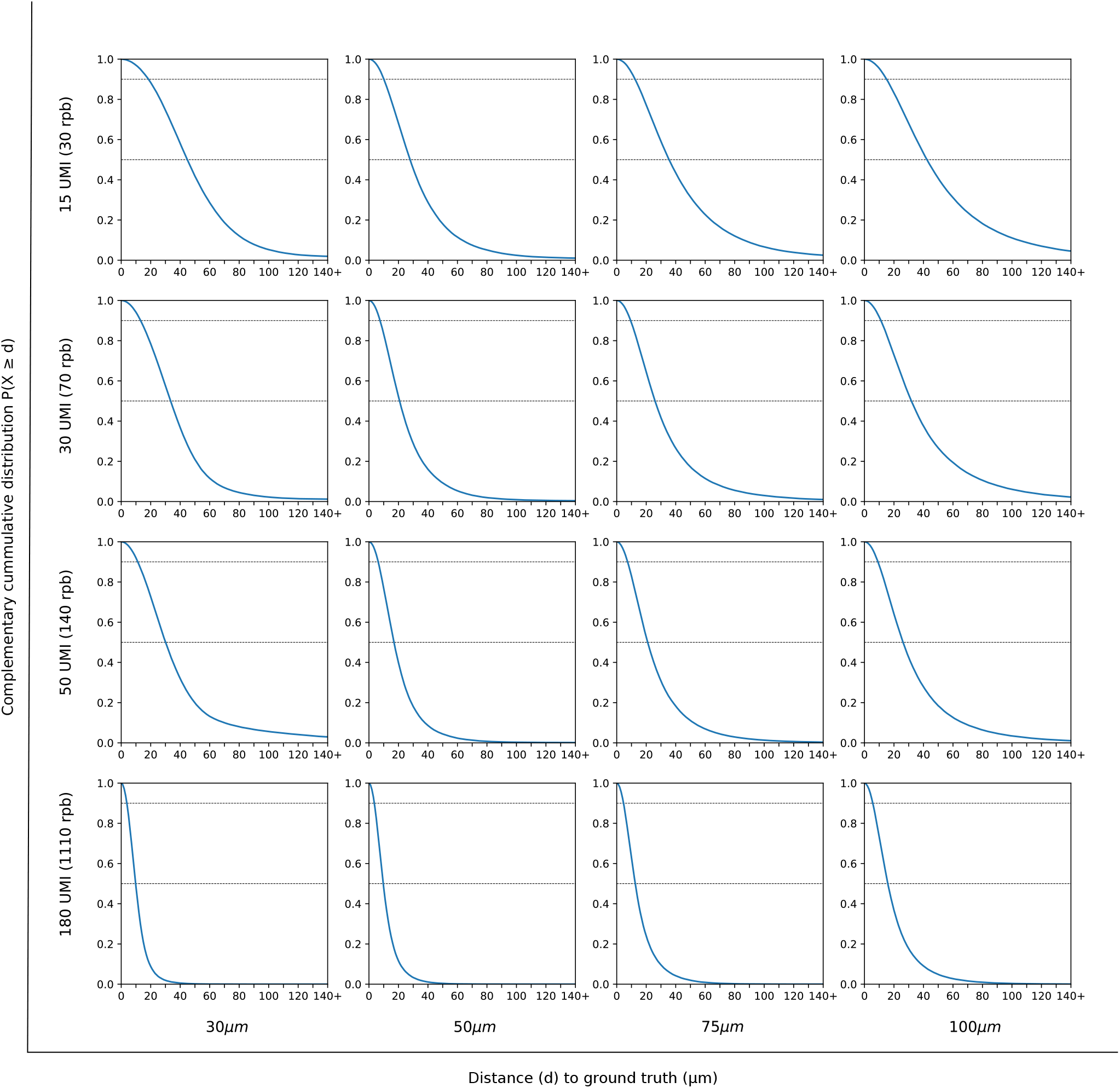
Proportion of beads beyond an alignment distance in *µm* for the simulations shown in Figure 4. Horizontal lines indicate the 50^th^ and 90^th^ percentiles. These distributions show that there are not a significant proportion of beads that cannot be placed in the reconstructions.

**Extended Data Fig. 6:**
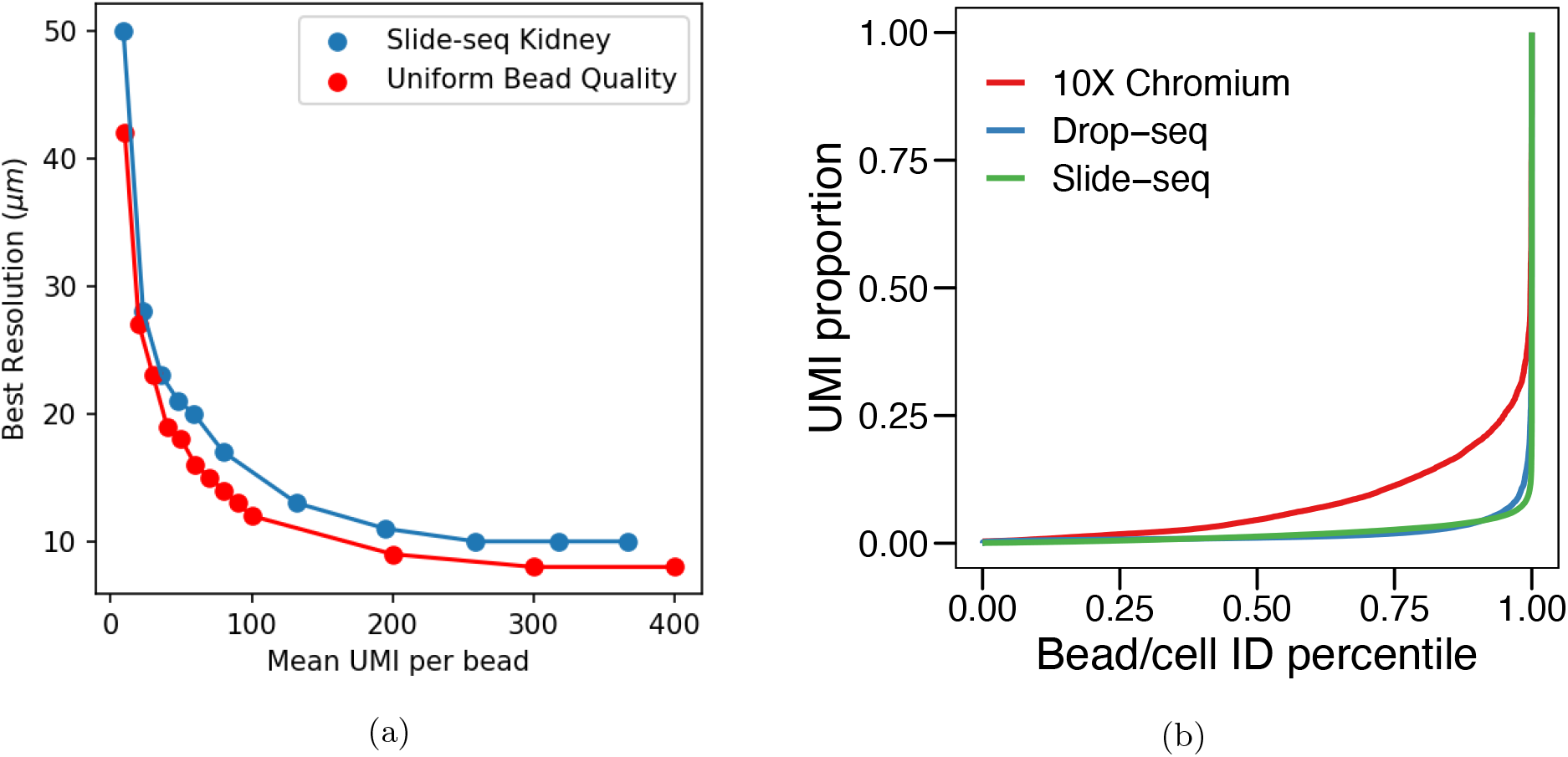
(a) Comparison of best reconstruction resolution for bead quality distributions from the Slide-seq kidney dataset and a distribution assuming uniform bead quality. As expected reconstructions using uniform bead quality outperform an experimental bead quality distribution. However, the performance is quite similar, supporting that the reconstructions are robust to changes in the bead quality distribution. (b) Comparison of the UMI distribution in the Slide-seq protocol compared to 10x and Drop-seq protocols. The less extreme distribution for the 10x protocol suggests that experimental bead quality distributions could be improved, leading to better reconstructions.

**Extended Data Fig. 7:**
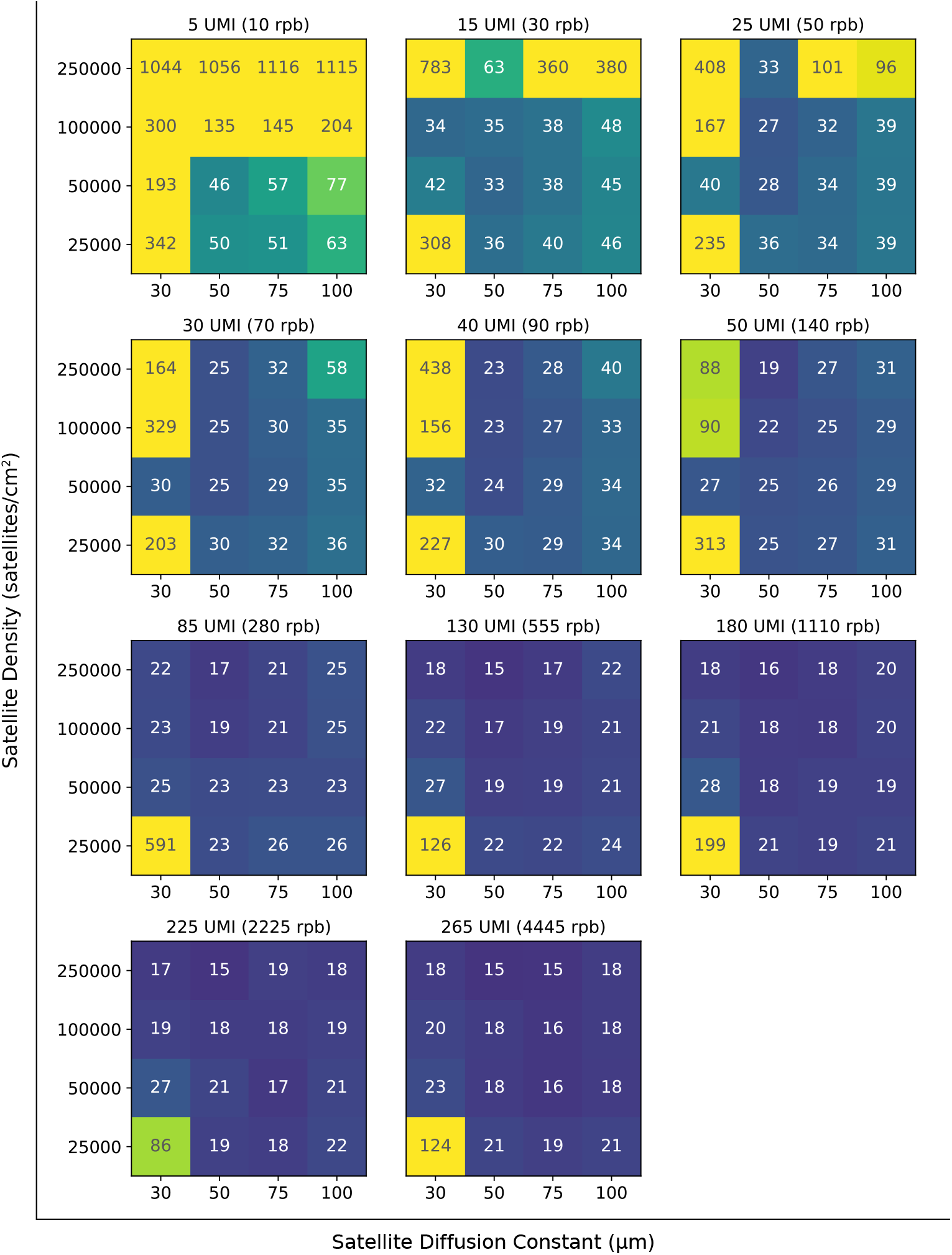
Summary of median alignment values in *µm* by number of satellites, diffusion level, and sequencing depth for reconstructions of bead positions from the Slide-seq kidney dataset. While the Slide-seq reconstructions performed slightly worse than the reconstructions on dense grids of beads, median alignment distances are still at the low end of the single cell range (10 - 100 *µm*) for a wide range of parameters. Further, these simulations show that sequencing depth can be adjusted to achieve a target resolution at the lowest sequencing cost.

**Extended Data Fig. 8:**
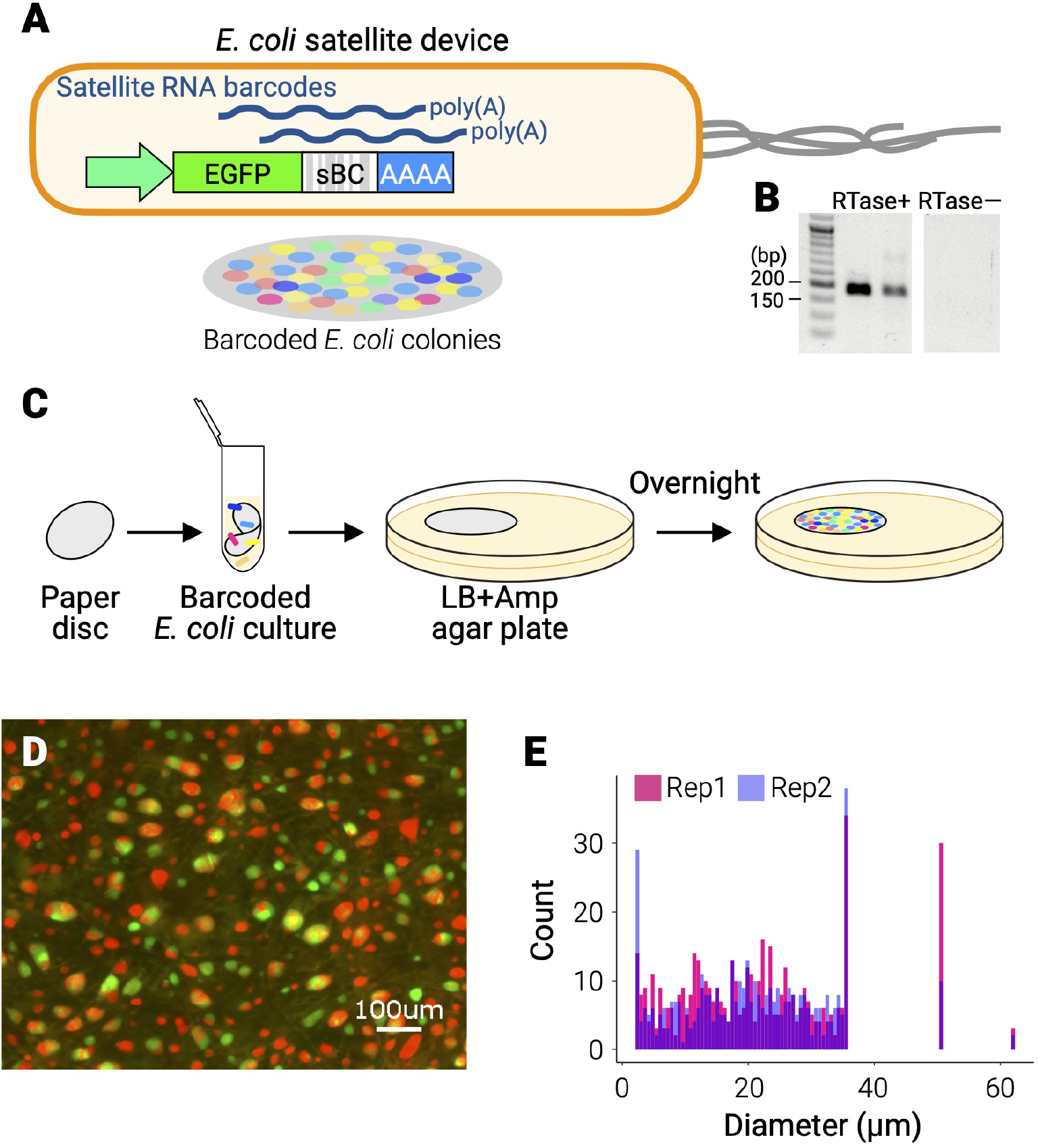
E. coli satellite device. (**A**) Design of the sBC expression unit. (**B**) Capturing of poly(A)ed sBC by reverse transcription using poly(T) RT primers. (**C**) Preparation of highly complex small sBC colonies. A paper disc is soaked in a complex E. coli cell culture, and placed on a solid agar plate for colony formation. (**D**) Preparation of small dense E. coli colonies. GFP- and RFP-expressing cells are mixed in 1:1 volume. (**E**) Diameter distribution of colonies formed on the paper disc.

## Supplementary Information

### Supplementary Equations

#### Mathematical model of the measurement process

To justify the use of manifold learning for spatial reconstructions, we show that the embedded beads belong to a two-dimensional manifold in satellite barcode space. We first present a general mathematical model for satellite diffusion. Second, we present a model for the sequencing process. Finally, we prove for the case where satellites are modelled by a point source that the embedded beads belong to a two-dimensional manifold in satellite barcode space.

#### Satellite diffusion model

We want to define a mathematical model for satellite diffusion. Let *X* be a compact subset of ℝ^2^, that defines our domain of interest e.g. [0, 1]^2^, or the unit disk. Let satellite *j* be a compact, connected set *S*_*j*_ ⊂ *X*. Notably, *S*_*j*_ can be a small region of the domain, or a single point, depending on the implementation of the satellite device (Figure 6). We treat each satellite as a source of satellite barcodes (sBCs). Each point in the domain *X* should receive signal (i.e. sBCs) from each satellite at some intensity.

We define *ψ*_*j*_ : *X* → ℝ_+_ as the signal intensity of satellite *j* over the domain. Intuitively, *ψ*_*j*_ should be a continuous, decreasing function of distance to satellite *j*. Let there be *M* satellite devices, we then define *ψ* as the vector of signal intensities received from each satellite at a given location on the domain:

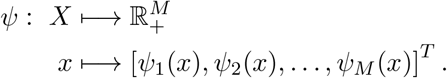

Physically, *ψ*(*x*) represents the abundance of satellite barcode molecules at position *x*. We refer to 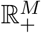 as the high-dimensional “satellite barcode space” (sBC space).

#### Sequencing produces a noisy embedding in sBC space

A collection of beads located at *x*_1_, …, *x*_*N*_ is embedded in sBC space as *ψ*(*x*_1_), …, *ψ*(*x*_*N*_) (Extended Data Fig. 1, black dots). The 2D surface of beads form a manifold ℳ in this high-dimensional sBC space (formally, ℳ is the image of *X* under *ψ*).

In practice, the sequencing process produces an *N* × *M* matrix **C** of counts, where each of the *N* beads has captured an integer number of unique barcode molecules from each of the *M* satellites. Each time a read is sequenced, a particular entry of this count matrix is incremented, starting from the all 0s matrix. This matrix thus provides a representation of the intensity distribution in terms of sBC counts, and the row vectors *c*_*i*_ of **C** are noisy measurements of the abundance of sBC molecules in the vicinity of each bead (Extended Data Fig. 1, red crosses).

We present two models for how the reads are sampled: a simple model and a more detailed one which accounts for variable bead qualities. The simplest possible model for how reads are sequenced is that each time a read is sampled, we get a read from satellite *j* and bead *i* with probability:

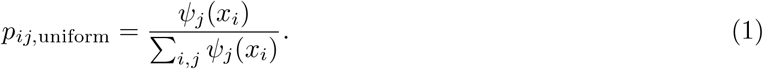

In this model, the probability of having a count between a bead and a satellite is based only on the signal intensity, and each bead would receive roughly an equal number of reads. However, we found that the Slide-seq data exhibited a wider range of counts per bead [4]. To account for this variation, we introduce the concept of bead quality, which is the proportion of reads captured by a bead relative to the total read count. Let *Q* be any distribution on (0, 1] (in practice, this can be generated from the distribution of counts in the Slide-seq data). Then, we can assign a quality to each bead as *q*_*i*_ ∼ *Q*. With this notion of bead quality, we can define the probability of getting a count between a bead and a satellite as:

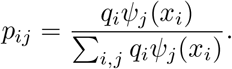

With these probabilities, we can model the sBC space with the matrix **C** of size *N* × *M*, capturing *R* samples from a multinomial distribution based on the signal intensities and bead qualities:

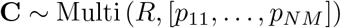

Because signal intensity decreases with distance, beads will only have non-zero barcode counts from local satellites, resulting in a sparse matrix **C**.

#### Barcode overlap induces a projection in sBC space

Another factor that is adding noise to the abundance of sBCs is the overlap of barcodes. While using a length *n* barcode there are *n*^4^ potential unique barcodes, and they do not occur uniformly in practice. Therefore, different satellites will have the same barcode, which makes recovering the bead positions more difficult. Mathematically, we model this by combining columns of **C** for satellites sharing a common barcode. This is equivalent to multiplying **C** by a projection operator **P**. The matrix we obtain from sequencing is therefore **Ĉ** = **PC**.

#### Distances in sBC space

Rows in **C** represent a sample from *p*_*i*_, which can be normalized to obtain a sample distribution. This normalization takes into account the varying bead qualities, as neighboring beads in the ground truth may have different bead qualities leading to large distances between their unnormalized rows in **C**. However, they will share similar normalized distributions. Alternately, the distance between two beads can be determined by performing a two sample test to quantify if the two samples come from a common distribution.

#### Manifold in sBC space

In this section we show that the embedded beads form a two-dimensional manifold in high-dimensional sBC space, rationalizing the use of manifold learning to recover their positions.

We consider the case where the satellites are point sources of sBCs, located at positions *y*_*j*_ ∈ ℝ^2^, *j* = 1, …, *M*. We then have *S*_*j*_ = {*y*_*j*_}. Next, we assume that sBCs diffuse according t o a radial point-spread function. Let *g* : ℝ _+_ → ℝ _+_, a decreasing continuous bijection, represent the radial decay of satellite signal intensity. Therefore, we have:

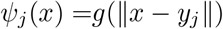

We now show that as soon as there are 3 non-aligned satellites, the beads embedded by *ψ* form a two dimensional manifold in sBC space:

##### Theorem 1.

*If M* ≥ 3 *and there exist j*_1_, *j*_2_, *j*_3_ ∈ [1, *M*] *such that* 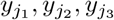 *are non-colinear, then ψ is a smooth embedding of X* ⊂ ℝ^2^ *into* 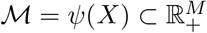, *which makes* ℳ *a smooth sub-manifold of* 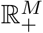 *as the image of ψ*.

*Proof. ψ* is a smooth embedding if it is a homemorphism onto its image. Therefore we need to prove that *ψ* is a continuous bijection from *X* to ℳ, with a continuous inverse.

Continuity of *ψ* is verified as the composition of continuous functions. By definition of ℳ as the image of *X* through *ψ*, we get that *ψ* is surjective.

Let us show that *ψ* is injective.

When *M* = 2, two different points in ℳ can have the same image through *ψ* since two circles respectively centered at *y*_1_, *y*_2_ with radii *g*^*−*1^(*ψ*_1_), *g*^*−*1^(*ψ*_2_) can intersect in at most two different places.

However, since *M* ≥ 3 and there exist *j*_1_, *j*_2_, *j*_3_ ∈ [1, *M*] such that 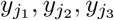 are non-colinear, then three circles with respective centers 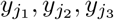 and radii 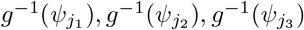 all intersect in at most one point, which means *ψ* is injective. Therefore, *ψ* is bijective.

Finally, the inverse *ψ*^*−*1^ is continuous since *X* is compact. Thus, *ψ* is a smooth embedding of *X* into ℳ, making ℳ a smooth sub-manifold of 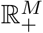.□

In practice we assume a Gaussian diffusion model, so we take *g* to be Gaussian (i.e. *g* : *x* → exp(− *x*^2^*/σ*^2^)).

Setting the width *σ* of this Gaussian then simulates different diffusion levels of the sBCs.

## Supplementary Discussion

Here we present further tests that demonstrate our method’s robustness to changes in physical parameters, bead quality, and bead positions.

### Robustness to physical parameters

As reported in the main text, we achieved mean alignment distances as low as 10*µm* and could achieve 20*µm* alignment distances for all but one combination of number of satellites and diffusion levels from 30 − 100*µm* (see Extended Data Fig. 3a). In these ranges, physical parameters do not limit whether a reconstruction is possible. Rather, some combinations result in better reconstructions for the same sequencing cost. While we achieved successful reconstructions down to 20*µm* of diffusion, they had high alignment errors even at the highest sequencing depth (see Extended Data Fig. 4). We did not achieve successful reconstructions for diffusion below 20*µm*. As preliminary experiments had already generated ∼ 20, 000 satellites*/*cm^2^, we did not test lower densities of satellites.

While the median alignment distance is a convenient statistic to compare between reconstructions, it is also important that there are not large number of outliers with high alignment distances, corresponding to beads that cannot be accurately placed. Extended Data Fig. 5 shows that in most cases *>* 90% of beads are no more than twice the median alignment distance.

### Robustness to bead quality

We repeated the simulations assuming uniform bead quality, i.e. we sampled reads according to *p*_*ij*,uniform_ from equation (1). As expected, the reconstructions improved compared to the experimental bead quality distribution. However, the improvements were moderate, with the best reconstruction only improving from 10 to 7*µm* and the best alignment error per UMI curves closely tracking (see Extended Data Fig. 6a). This suggests the method is robust to changes in the distribution of bead qualities. Further, we believe the experimentally obtained bead quality distribution is a conservative estimate of what could be achieved in practice. For example, the 10x protocol achieves a less extreme distribution than Slide-seq and Drop-seq which employ similar library preparation protocols (See Extended Data Fig. 6b).

### Slide-seq Reconstructions

Using bead positions from the Slide-seq kidney, cerebellum 2, and hippocampus datasets we were able to achieve alignment distances as low as 15*µm* and could achieve 30*µm* alignment distances for all but one combination of number of satellites and diffusion levels from 30 − 100*µm* (See Extended Data Fig. 7 and Extended Data Fig. 3b). We believe that the reconstructions were slightly worse for the Slide-seq bead positions because only beads positioned over the tissue were included in the dataset, leading to regions without beads. UMAP assumes that sampled points are uniformly distributed, so our reconstructions perform best when beads have close to uniform density. In practice, all beads would be sequenced leading to roughly uniform density and making these tests a conservative estimate of reconstruction quality. Despite this limitation, the reconstructions with the Slide-seq positions were still at the low end of the single-cell range.

